# Evaluating the Identifiability of Temporal Delay and Response Attenuation in Endometrial Decidualization

**DOI:** 10.64898/2026.11.03.764027

**Authors:** Elena Krikun, Wael Fahmy, Abedalrhman Alkhateeb

**Affiliations:** Lakehead University Thunder Bay, Ontario, Canada

## Abstract

Endometriosis is associated with impaired endometrial decidualization, yet conventional differential expression analyses cannot readily distinguish delayed transcriptional activation from reduced response amplitude. These mechanisms may produce similar expression differences at individual time points while implying distinct biological interpretations. Establishing when they can be distinguished is therefore a computational identifiability problem.

We implement an identifiability-aware analysis of temporal displacement and amplitude attenuation in longitudinal transcriptomic responses. The approach combines reference trajectories, constrained amplitude-only and amplitude-plus-delay models, analytical examples of observational equivalence, and simulation-based assessments of mechanism distinguishability. Held-out time-point evaluation and culture-level sensitivity analyses examine the stability of model-dependent conclusions. In GSE246591, a candidate-delay-dependent dynamic-range filter excluded 30.9–38.3% of initially dynamic features under a hypothetical 24-hour shift across four endometrial stromal cultures. A two-component diagnostic separating retained dynamic range from affine shape similarity revealed that small projection residuals can arise from either loss of observable dynamics or similarity of response shapes. These findings demonstrate why a single separation statistic may be insufficient for interpreting temporal identifiability. Complementary independent in vivo analyses provided biological context. In GSE51981, an endometrial progesterone-response-associated transcriptional score was reduced in endometriosis samples (phase-model estimate 0.155, 95% CI 0.222 to 0.088). GSE6364 showed a directionally consistent but statistically inconclusive difference (0.033, 95% CI 0.068 to 0.002). Neither observation independently establishes a temporal delay.

Together, these results support selection-aware evaluation of temporal transcriptomic responses, distinguishing structural ambiguity and loss of dynamic signal from evidence of a biological timing change.

## 1 Introduction

Decidualization is the hormone-associated differentiation of human endometrial stromal cells that prepares the uterine lining for implantation [1]. An attenuated decidualization-associated transcriptional signal in endometriosis has been reported in multiple endometrial studies [2, 3]. Yet a lower signal at one sampling time can arise from a smaller response, a later response, altered trajectory shape, menstrual-cycle composition, or an unmeasured cellular mixture. Distinguishing those explanations requires information beyond cross-sectional differential expression. Menstrual-cycle-associated confounding further complicates endometrial comparisons [4].

Temporal gene-expression methods answer related but distinct questions. *DEtime* estimates when two perturbation-specific time courses diverge using Gaussian processes [5]; *ImpulseDE2* tests and models time-dependent expression using an impulse family [6]. Neither method, without a suitable matched design and specified nuisance parameters, establishes that a lower single-time signal is specifically a delayed version of another response. Identifiability analysis asks whether different parameter settings can generate the same observable data, a question with a long history in dynamical modeling [7, 8].

We operationalize the distinction in two steps. First, we identify when a shifted curve can be represented exactly by changing only its amplitude and baseline (*observational aliasing*). Second, we quantify the remaining separation on a verified endometrial RNA-sequencing schedule, with explicit checks of expression assayability, reference-curve assumptions, and uncertainty limits. The resulting audit separates algebraic nonidentifiability from what measured expression values can establish. A supplementary software example documents a related computational workflow, but does not serve as biological validation.

## 2 What does it mean to identify a temporal delay?

### 2.1 A minimal model and an exact criterion

For sampling times *T* = {*t*_1_, …, *t*_*m*_*}* and a specified reference expression profile *f* (*t*), compare an amplitude-only response with an amplitude-plus-shift response:

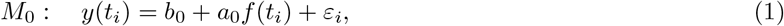

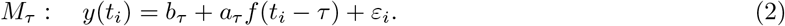

Here *b* is the baseline expression level, *a* is the response amplitude, and *τ* is a displacement relative to a chosen reference trajectory, *not* an experimentally observed molecular-onset time. The *ε*_*i*_ represent measurement error or departures from the assumed trajectory. We first consider the noise-free case with unconstrained baseline and nonzero signed amplitudes.

#### Proposition 1

(Observational equivalence on a finite time grid). *Let* 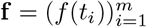 *and* 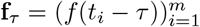 .

*The two response families can generate the same noiseless vector with nonzero amplitudes if and only if*

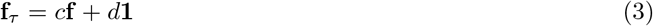

*for an admissible c* ≠ 0 *and offset d. If amplitudes are restricted, c must also respect those restrictions*.

*Proof*. Equality *b*_*τ*_ **1** + *a*_*τ*_ **f**_*τ*_ = *b*_0_**1** + *a*_0_**f** yields *c* = *a*_0_*/a*_*τ*_ and *d* = (*b*_0_ −*b*_*τ*_ )*/a*_*τ*_ . Conversely, substituting Eq. (3) and setting *a*_0_ = *a*_*τ*_ *c, b*_0_ = *b*_*τ*_ + *a*_*τ*_ *d* makes both vectors identical. Constant or zero-amplitude edge cases must be treated separately. □

For example, *f* (*t*) = *e*^*kt*^ has *f* (*t* −*τ* ) = *e*^*−kτ*^ *f* (*t*), whereas *f* (*t*) = *αt* + *β* has *f* (*t*− *τ* ) = *f* (*t*) −*ατ* . For a *fixed* shift *τ*_0_, *f* (*t*) = *e*^*kt*^*q*(*t*) with *q*(*t*− *τ*_0_) = *q*(*t*) also aliases. These are familiar algebraic counterexamples, not new solution families. Such exact identities assume the shape extends over the shifted domain: a separate rule that clamps *f* (*t*) before stimulation may break that identity.

### 2.2 A measurable, but conditional, diagnostic

Set *X* = [**1, f** ], let *P*_*X*_ = *XX*^+^ be the least-squares projector (with *X*^+^ the Moore–Penrose pseudoinverse), and define

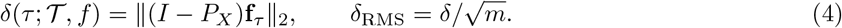

#### Proposition 2

(Signal–shape decomposition of a projection residual). *Let* **g** = **f**_*τ*_ *be the modeled shifted curve and suppose* range(**f** ) *>* 0. *Define D* = *δ*_RMS_*/* range(**f** ), *R* = range(**g**)*/* range(**f** ), *and, for nonconstant*

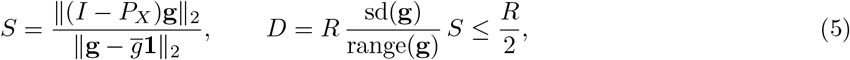

*where* sd(**g**) *uses divisor m. If* **g** *is constant, D* = 0, *R* = 0, *and S is undefined. For two varying curves, the unconstrained signed-amplitude projection gives S*^2^ = 1 − *r*(**f**, **g**)^2^, *where r is their Pearson correlation*.

*Proof*. Because **1** ∈ col(*X*), the projection residual of **g** is no larger than the centered norm of **g**. With its range *B*, sd(**g**) ≤ *B/*2 by the range–variance inequality. Substitution yields Eq. (5). The correlation identity follows from ordinary least-squares regression of one centered vector on the other.

A small *D* can therefore arise from weak retained signal (*R* small) or similarity after affine reparameterization (*S* small). These are different limits on interpretation. Neither quantity is a confidence interval or a calibrated onset test. The two components do not alone determine *D*; its exact product also involves sd(**g**)*/* range(**g**).

#### Which quantities remain identifiable?

For the illustrative exponential reference *f* (*t*; *k*) = *e*^*kt*^, a delayed response *y*(*t*) = *b* + *ae*^*k*(*t−τ*)^ can be written as *b* + *Ae*^*kt*^ with the *identifiable combination A* = *ae*^*−kτ*^ . No expression-only observation separates *a* from *τ* in this unrestricted family. Yet, in noise-free observations at three equally spaced times *t, t* + *h, t* + 2*h*, a nonzero response with *k* ≠ 0 obeys

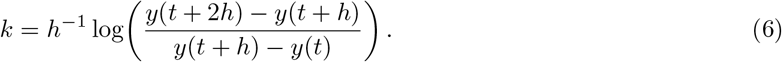

Thus *k* may remain identifiable even when *τ* does not. This is a model-specific illustrative reparameterization, not a general estimator of a biological rate; near-flat curves or noisy observations can make *k* practically unrecoverable [9].

#### Identifiability-aware feature eligibility

If Eq. (3) holds on the sampled grid, the identity

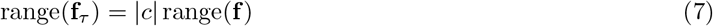

is exact. Therefore requiring *both* observed and modelled-shifted ranges to exceed a fixed threshold can exclude some exactly aliased trajectories in a shift-dependent way. This conclusion concerns a candidate-dependent *shifted-range* gate; it does not establish that conventional filtering on observed expression alone invalidates temporal analyses. The direction of the selection effect depends on the trajectory, shift and pre-onset boundary rule [10]. We report eligibility separately from projection residuals.

This is the residual after allowing the unshifted curve to change its baseline and *signed* amplitude. Zero residual denotes exact equivalence on the sampled grid; nonzero residual indicates geometric separation *conditional on the assumed reference and boundary rule*. It is neither a p-value nor a confidence interval and cannot determine an onset time by itself. With only two sampled times and unrestricted offset/amplitude, even distinct shifts can always have zero residual when *X* spans the observation space. Sampling more times can break some ambiguities but cannot resolve a globally exponential or affine identity by itself.

## 3 Datasets and computational protocol

### 3.1 Observed endometrial time course

The Gene Expression Omnibus (GEO) dataset **GSE246591** contains RNA-sequencing (RNA-seq) profiles of four primary human endometrial stromal-cell cultures, S169, S170, S506, and S508, each sampled at **0, 3, 6**,**9, 12, 18, 24, 36, 48, and 96 hours** after decidualization induction [11]. We verified all 40 culture-by-hour columns in the original deposited counts file (SHA-256 in Supplementary Methods). There are 60,676 source feature rows. These are source identifiers, not necessarily 60,676 uniquely mapped genes. No independent within-culture, same-hour replicate series is available for estimating measurement variance or ground-truth delay.

For each feature and culture, raw counts were divided by the total reads assigned to the sample, scaled to one million (counts per million; CPM), and transformed as log_2_(1 + CPM). We considered hypothetical displacements *τ* ∈ { 3, 6, 12, 24} hours. The measured ten-point curve was connected by *piecewise-linear interpolation*; for shifted arguments below time zero we imposed *f* (*t <* 0) = *f* (0). No expression at an unmeasured shifted time was treated as a new observation. For each candidate displacement we computed Eq. (4) and divided *δ*_RMS_ by the measured ten-point expression range. Features entered each descriptive summary only if both their measured and modeled shifted expression ranges were at least 0.5 log_2_(1+CPM). Consequently, the set of included features changes with shift and culture; our summaries are not paired gene-level contrasts.

### 3.2 Evidence roles and safeguards

Sample identities, input checksums, eligibility rules and model-specific statistics are recorded in the Supplement. We present no estimate of power or classification accuracy for molecular onset. The threshold of normalized projection residual below 0.05 is *descriptive only*: it is not derived from the experimental error distribution. We separately describe a retrospective shape-sensitivity exercise on 48 features but do not call it a head-to-head benchmark with DEtime or ImpulseDE2.

**Table 1.** Distinct analytical roles. The reported datasets are *not pooled*; samples do not represent exchangeable units across studies.

| GEO series | Design | Role / boundary |
|---|---|---|
| GSE246591 | Four cultures; ten sampling times each | Reference-shift geometry; no known biological lag or within-time replicate variance. |
| GSE94036 | PGR/FOXO1 perturbation in stromal cells | Source of the PGR-oriented score, not a timed-lag experiment. |
| GSE51981 | Tissue microarrays; MSE subset 28 vs 8 | Fixed-score clinical contrast, not measured delay. |
| GSE6364 | Tissue microarrays; MSE subset 9 vs 8 | External directional score check, not an hourly time course. |

## 4 Results

### 4.1 Exact counterexamples: why timing alone may be unidentifiable

All three algebraic examples in Fig. 2 satisfy Eq. (3) for the specified shifts and allowed nuisance parameters. Thus zero-noise expression values alone need not distinguish a temporal shift from rescaling or changing background expression. These mathematical witnesses do not establish that the corresponding shape is biologically correct. A separate historical implementation verified the three identities to numerical tolerance; we treat the identities themselves, rather than an inferred biological lag, as the result.

**Figure 1:**
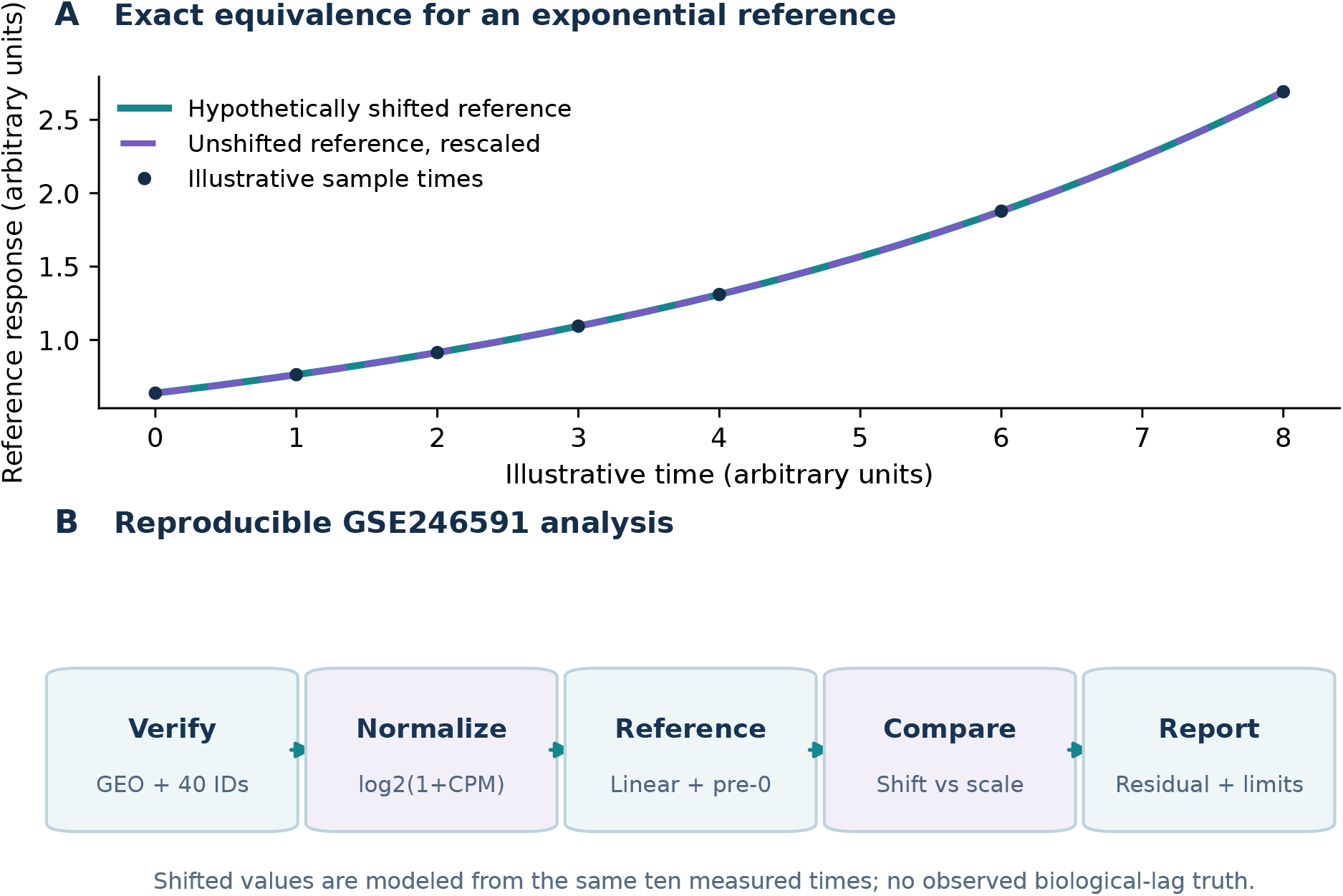
Exact ambiguity and analytical pipeline. (A) The two profiles overlap algebraically for an illustrative exponential reference; dots are *not* measured RNA-seq. (B) The mandatory audit sequence checks sample identities, fixes the reference and nuisance parameters, compares amplitude with displacement, assesses remaining geometric separation, and reports limits. The programmatic implementation records source files and checksums.

**Figure 2:**
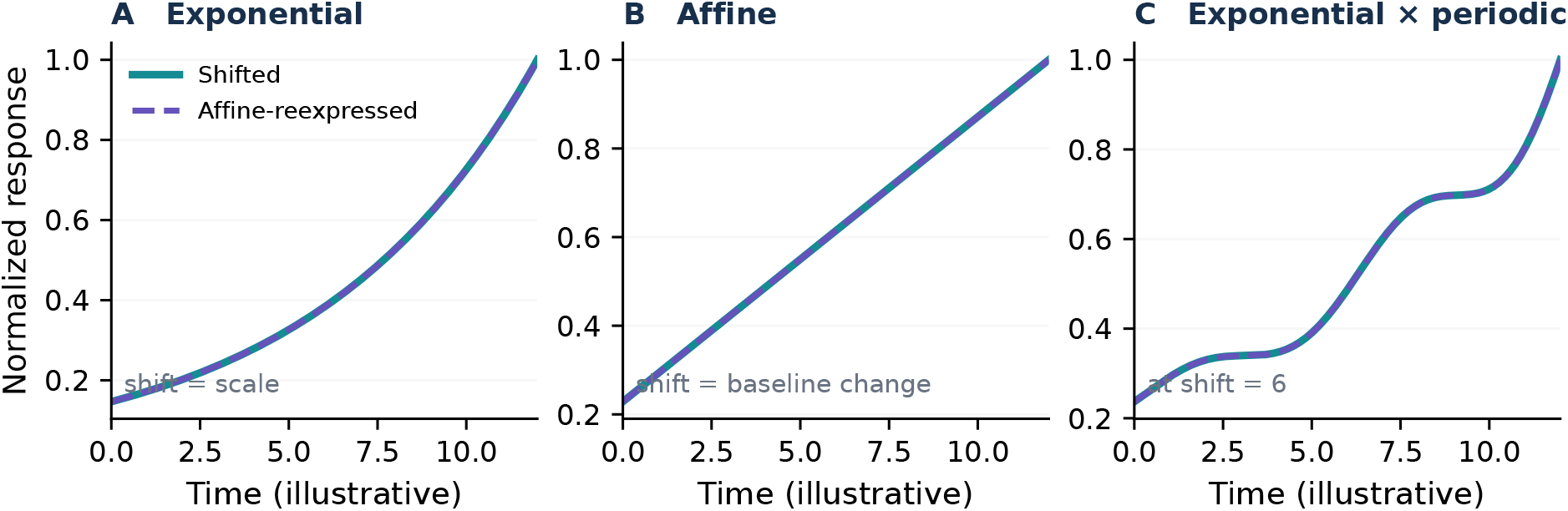
Exact analytic counterexamples, not measured data. An exponential, an affine function and an exponential multiplied by a periodic factor can each alias a specified shift through scale and/or offset. The periodic construction establishes equality for its stated shift, not all possible shifts. The curves here are unclamped; the empirical analysis uses an explicit flat pre-zero boundary.

### 4.2 Real measured trajectories show conditional, heterogeneous separation

The deposited GSE246591 counts matrix passed the 40/40 sample-identity and integer-count checks. Across the four cultures and four shifts we obtained 16 descriptive summaries (Supplementary Table S1). For *τ* = 3 hours, median *δ*_RMS_*/*range(*f* ) was 0.220 for S169 and 0.153–0.165 for the remaining cultures. At 6 hours the four medians were 0.206–0.214; at 12 hours they were 0.226–0.231; at 24 hours they were 0.209–0.219 (Fig. 3). Thus a general assertion that *most* measured response curves are nearly exactly compatible with a shifted-and-rescaled response is not supported by this diagnostic.

**Figure 3:**
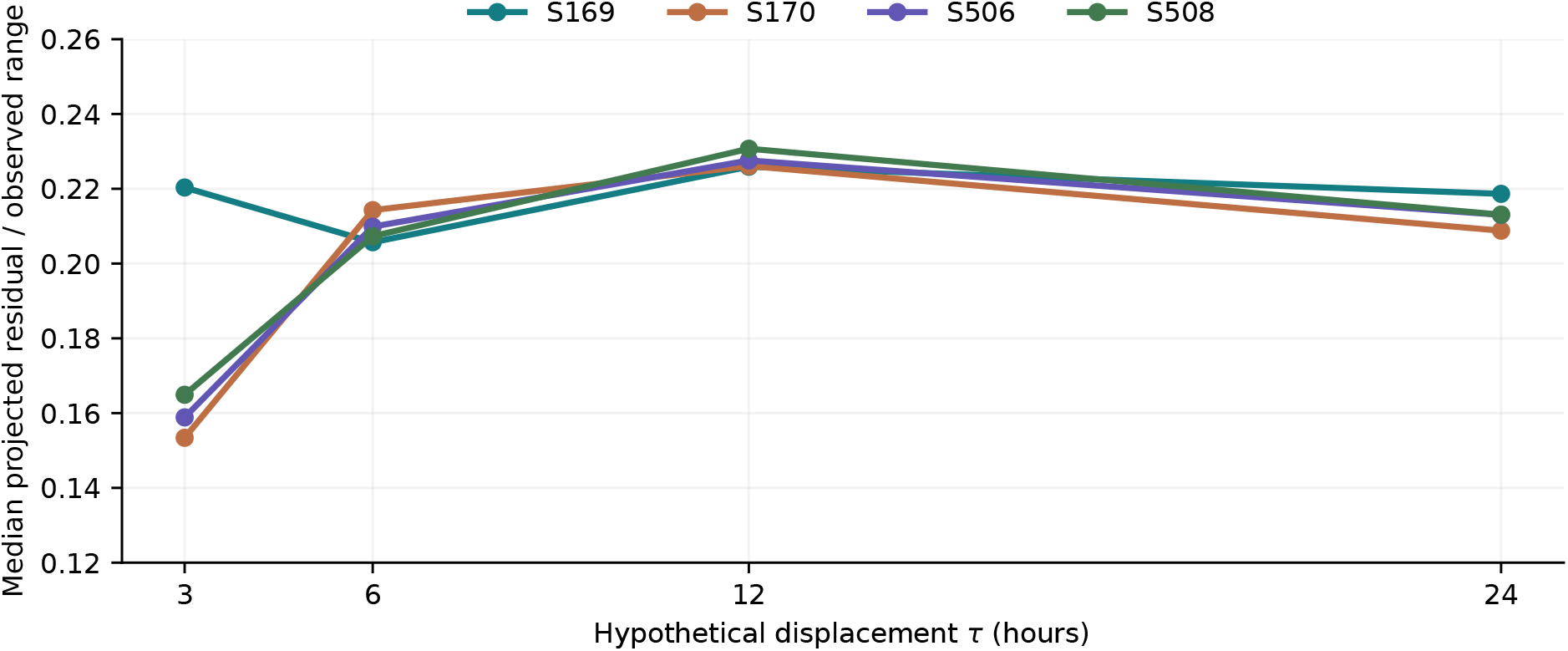
Measured-source, model-conditional GSE246591 geometry. Medians over included feature rows of the root-mean-square residual after projecting a hypothetical displaced curve onto baseline and signed amplitude, divided by each feature’s measured expression range. The values are computed from actual GEO counts, while displaced coordinates rely on piecewise-linear interpolation and the flat pre-zero rule. Feature eligibility differs by *τ* (Supplementary Fig. S3); connecting lines is for readability, not evidence of a continuously estimated lag.

At the descriptive cutoff *δ*_RMS_*/*range(*f* ) *<* 0.05, the fraction of included features ranged from approximately 0.02% to 3.20% across the 16 culture-by-shift settings. This is neither an estimate of the proportion of genuinely delayed genes nor a noise-calibrated test: the source includes only one profile per culture-hour and the hypothetical shifted values are computed from the same measured series. Larger shifts also change the eligible feature universe; for example S169 included 14,019 rows at three hours but 9,116 at 24 hours. We therefore do not interpret across-shift median differences as paired gene-level effects. The reported proportions also condition on *both* dynamic-range gates. In particular, exact aliased profiles with shrinking modeled response ranges need not remain eligible (Eq. (7)). A selection-aware audit instead reports the denominator passing the observed-range gate, the additional exclusions caused by the candidate shift, and the distribution of the same geometric residual in retained and excluded features (Supplementary Methods). This is a retrospective diagnostic, not a population estimate of lag identifiability.

### 4.3 Separating signal loss from affine shape similarity

The two-component audit of the same GSE246591 matrix identifies distinct explanations for a small *D* (Fig. 4). Among features passing the initial observed-curve range screen, 982 culture–feature records at a three-hour candidate shift had *D <* 0.05; 977 (99.5%) also had *S <* 0.20, whereas none had *R <* 0.20. Within each culture the near-zero group’s median *R* was 0.968–0.975 and median *S* was 0.131–0.141. Thus short-shift small residuals commonly arose from affine shape similarity despite retained dynamic range, under the specified reference model.

**Figure 4:**
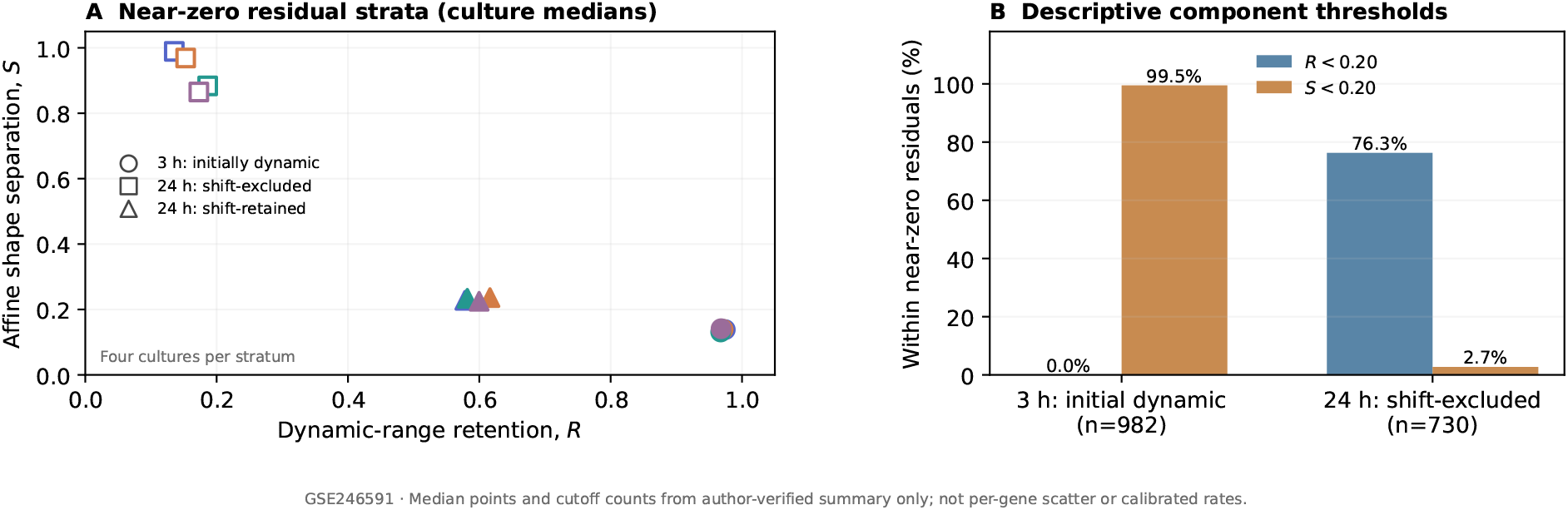
Two explanations for a small geometric projection residual. (A) Each point is a *culture-level median among near-zero* (*D <* 0.05) records, not a per-gene joint scatter; the three symbols distinguish the three-hour initially dynamic stratum from 24-hour shifted-range-excluded and retained strata.(B) Descriptive percentages below *R* = 0.20 and *S* = 0.20 within the corresponding near-zero groups. The groups differ in size and selection rule; thresholds are illustrative, not calibrated tests. Source: GSE246591 and the archived author-run dual-component audit.

At a 24-hour shift, 730 culture–feature records were both excluded by the shifted-range screen and below *D <* 0.05. Of these, 557 (76.3%) had *R <* 0.20, while only 20 (2.7%) had *S <* 0.20; one shifted curve was exactly flat and had undefined *S*. Median *R* among these low-*D* excluded features was 0.136–0.186 across cultures, whereas median defined *S* was 0.866–0.989. Their small residuals more often reflected loss of the shifted signal than similarity of two varying shapes. The 122 near-zero records that remained shift-eligible at 24 hours showed comparatively higher median *R* (0.578–0.616) and lower *S* (0.225–0.236). The illustrative 0.20 cutoffs do not define statistical significance.

The distinction also explains why the secondary dynamic-range gate changes the denominator: of 56,988 initially dynamic culture–feature records at 24 hours, 19,676 (34.5%) were excluded by the shifted-range condition. Among all initially eligible records, 852 (1.50%) had *D <* 0.05, compared with 122/37,312 (0.33%) of the retained set. We do *not* interpret this difference as proof that filtering preferentially removes structurally aliased trajectories: Eq. (5) makes small *D* mechanically possible when *R* is small. A fixed intersection of eligible features across all four candidate shifts (7,858–9,960 rows per culture) provides a stable denominator for supplementary descriptive comparisons. Additional culture- and shift-specific summaries are reported in the Supplement.

### 4.4 Sensitivity to the assumed reference-curve shape

A separate, retrospectively selected *fixed* 48-feature subset was evaluated using piecewise-linear interpolation, a Gaussian-process (GP) approximation with fixed parameters, and a two-sigmoid impulse-like approximation. In leave-one-time-point-out reconstruction, linear interpolation had the lowest median feature-level root-mean-square error (RMSE) in each culture (0.595–0.737 log_2_CPM). The GP and impulse-like curves often produced smaller conditional residuals for a three-hour hypothetical shift, illustrating the sensitivity of Eq. (4) to the reference representation. The pilot emitted numerical warnings from matrix operations despite finite exported tables. We therefore classify this exercise as *exploratory and not numerical method validation*; its complete descriptive readouts and caveats are in Supplementary Table S3. These approximations are not the published implementations of DEtime or ImpulseDE2; they serve only to test sensitivity to curve representation.

### 4.5 Independent biological context does not establish temporal delay

The PGR (progesterone receptor)-dependency score was derived from a separate stromal-cell perturbation experiment (GSE94036) and evaluated in endometrial cohorts [2, 3, 12–14]. In the 28 endometriosis versus eight strict-control mid-secretory endometrium (MSE) samples from GSE51981, the estimated score difference was −0.154922. The *phase-interaction model* using all eligible menstrual phases (*n* = 109) yielded a 95% confidence interval (CI) based on HC3 heteroskedasticity-robust standard errors [−0.222227, −0.087617] for the MSE contrast and *p* = 1.386 ×10^*−*5^. A separate mid-secretory-only model with HC3 standard errors produced [−0.223889, −0.085955] (*p* = 6.247 ×10^*−*5^); adjusting that MSE-only model for three composition-proxy components gave −0.076561 (95% CI [−0.133443, −0.019678], *p* = 0.009975). These are different models and must not be assigned one another’s p-values.

In the independent GSE6364 mid-secretory subset (nine endometriosis cases and eight controls), the frozen-score difference was −0.032765. Its unequal-variance *Welch* 95% CI was [−0.067656, +0.002126], with *p* = 0.063752. Its direction was consistent but its interval included zero. The GSE51981 values from the phase model and the GSE6364 Welch estimate explain the rounding in the registered abstract. Neither endometrial comparison repeatedly samples the *same women* over hours after a controlled perturbation; no delayed molecular onset can be inferred from them. We do not pool effect sizes across these distinct estimands.

## 5 Discussion

The central distinction is between *structural ambiguity* (different mechanisms giving exactly the same noiseless observations) and *empirical information* (the separation suggested by an observed curve under specified assumptions). Algebra shows that some shape families can alias delay exactly. With an empirical curve, projection residuals quantify the distance between hypothetical shifted and unshifted profiles only after reference interpolation, boundary behavior, background and amplitude freedom have been specified. Even a nonzero residual cannot imply practical distinguishability without replicate-level noise and appropriate dependence modeling [7, 8]. The actual GSE246591 grid showed heterogeneous, mostly non-negligible normalized distances among the eligible features rather than pervasive near-zero residuals. That descriptive finding refines the motivating question; it does not recover the onset of a biological event.

A constructive positive implication is that when a shift is absorbed by an amplitude–baseline transformation, a parameter describing *shape* may remain measurable. For an exponential family, the rate *k* is an example (Eq. (6)); time to a specified observed plateau is not automatically an identifiable substitute and would require its own model-specific analysis. The appropriate estimand therefore depends on which parameters survive the nuisance reparameterization.

The signal–shape decomposition additionally shows why a low range-normalized residual is not, by itself, evidence of affine equivalence: reduced shifted dynamic range can lower the residual without reducing the relative shape-separation ratio. We recommend reporting signal retention, affine shape separation, non-evaluable flat profiles and eligibility counts together, rather than using a single near-zero residual fraction. This is an interpretive auditing protocol built from standard regression geometry, not a novel statistical test or a proven method for recovering onset.

The main limits are a single measured expression vector per culture-hour, no known imposed lag, no matched untreated curve at every time, uncertain interpolation and pre-zero behavior, source feature identifiers not mapped to unique genes, and retrospective eligibility/filter choices. Thousands of feature rows should not be counted as thousands of independent biological replicates. In addition, the amplitude projection is signed; a constrained nonnegative-amplitude model would yield different distances when fitted slopes are negative. The Supplement distinguishes that issue and the separate fixed-panel historical analyses. The curve-shape pilot is not a validated comparison with published state-of-the-art methods and emitted numerical warnings; its values must not be used to rank algorithms.

Related preprints by the present authors discuss assayability and cohort transfer [15], and the distinction between computational stability and biological interpretation [16]. They motivate separate reporting of measurement compatibility, statistical reproducibility and biological meaning, but do not validate this endometrial timing hypothesis. A decisive follow-up would preregister amplitude and curve-shape constraints, collect matched untreated time courses from independent donors with within-time biological replicates, impose a known perturbation delay, and assess candidate estimators on the same prediction target. Such a study would also permit a fair comparison with DEtime, ImpulseDE2 and other established time-course approaches. Where those conditions are absent, the correct result is a limit on interpretability, not a claimed delayed response.

## 6 Conclusions and software availability

Delay and attenuation can be observationally equivalent even with perfect measurements, but how closely they agree on a real sampling grid is reference- and assumption-dependent. Our endometrial RNA-seq audit provides reproducible *descriptive* evidence for that distinction without claiming biological-onset recovery or a new identification theorem. Code and reproducibility instructions: https://github.com/Helen2813/paper7. The separate OnsetAudit application, its genuine screenshot and its mouse-cell elapsed-exposure-time benchmark are documented in the Supplement; they are not evidence for human endometrial lag.

## Notes

https://www.ncbi.nlm.nih.gov/geo/query/acc.cgi?acc=GSE246591

https://www.ncbi.nlm.nih.gov/geo/query/acc.cgi?acc=GSE51981

https://www.ncbi.nlm.nih.gov/geo/query/acc.cgi?acc=GSE6364

https://www.ncbi.nlm.nih.gov/geo/query/acc.cgi?acc=GSE94036

https://github.com/Helen2813/paper7

### Competing Interest Statement

The authors have declared no competing interest.

