## Supplementary material for "Evaluating the Identifiability of Temporal Delay and Response Attenuation in Endometrial Decidualization": suplimentary

### S1. Explicit definitions and reproducibility

**A sampling grid** is the set of times at which actual RNA is measured. **A reference trajectory** $f(t)$ is an assumed function describing expected expression over that grid; measurements alone do not determine the function between points. **An onset** is the biological time a molecular process begins, which is not directly labeled in these studies. **A temporal displacement** $\tau$ is a hypothetical shift of a specified reference. **Response attenuation** is a change in amplitude at fixed timing. **Structural nonidentifiability** is exact observational equivalence in the assumed model even without noise. **Practical identifiability** additionally requires an error model and sufficient signal relative to that error. A **negative control** is an explicitly specified comparator expected not to carry a target effect, not merely any unused sample.

The public repository is `https://github.com/Helen2813/paper7`. The present paper focuses on the GSE246591 depositor count matrix, not the distinct day-scale study GSE243039. Source SHA-256 for the GSE246591 count file is `1d340b52b0eac564f73fe892ab28560748a122a7d3d51e21aa3c13e8a667cb08`. This checksum was reported by the verified Mac execution with 40/40 exact culture-time sample matches (0, 3, 6, 9, 12, 18, 24, 36, 48, 96 hours). The original first column is labeled `Ensemvl.ID`; we preserve the source identifier and do not silently assert one-to-one gene-symbol mapping. The 60,676 features may contain low-expression features and are not independent replicate observations.

Expression processing: let $C_{gs}$ be the raw integer count for feature $g$ and sample $s$, and $L_s = \sum_g C_{gs}$ its library size. We set $f_{gs} = \log_2(1 + 10^6 C_{gs}/L_s)$. Each culture has a ten-point $f_g(t_i)$. The modeled shifted value is $\widetilde{f}_g(\max(0, t_i - \tau))$ from linear interpolation among measured knots, with values before zero clamped to the initial level. This is a model assumption, not an additional measurement. Least squares profiles out a free background $b$ and a signed amplitude $a$:

$$\delta_{g,\mathrm{RMS}}(\tau) = \frac{1}{\sqrt{10}} \min_{a,b\in\mathbb{R}} \left\| \widetilde{\mathbf{f}}_{g,\tau} - b\mathbf{1} - a\mathbf{f}_g \right\|_2.$$

Our descriptive normalized diagnostic is $\delta_{g,\mathrm{RMS}}/\mathrm{range}(\mathbf{f}_g)$. We include rows where *both* measured and modeled shifted expression ranges are $\geq 0.5 \log_2(1+\mathrm{CPM})$. The feature set is therefore shift-dependent. The arbitrary 0.05 cutoff defines the table's small-residual fraction; it is not based on donor-level variance, and the counts of features should not be interpreted as independent sample sizes. The underlying input produced by the user is reproducible with the bundled `scripts/compute_gse246591_delta.py`, which strictly checks the expected 40-column mapping and source checksum. The input was processed locally by the study author; this Overleaf distribution includes its 16 console summaries but not the original full count matrix.

#### S2. Software: the actual OnsetAudit application

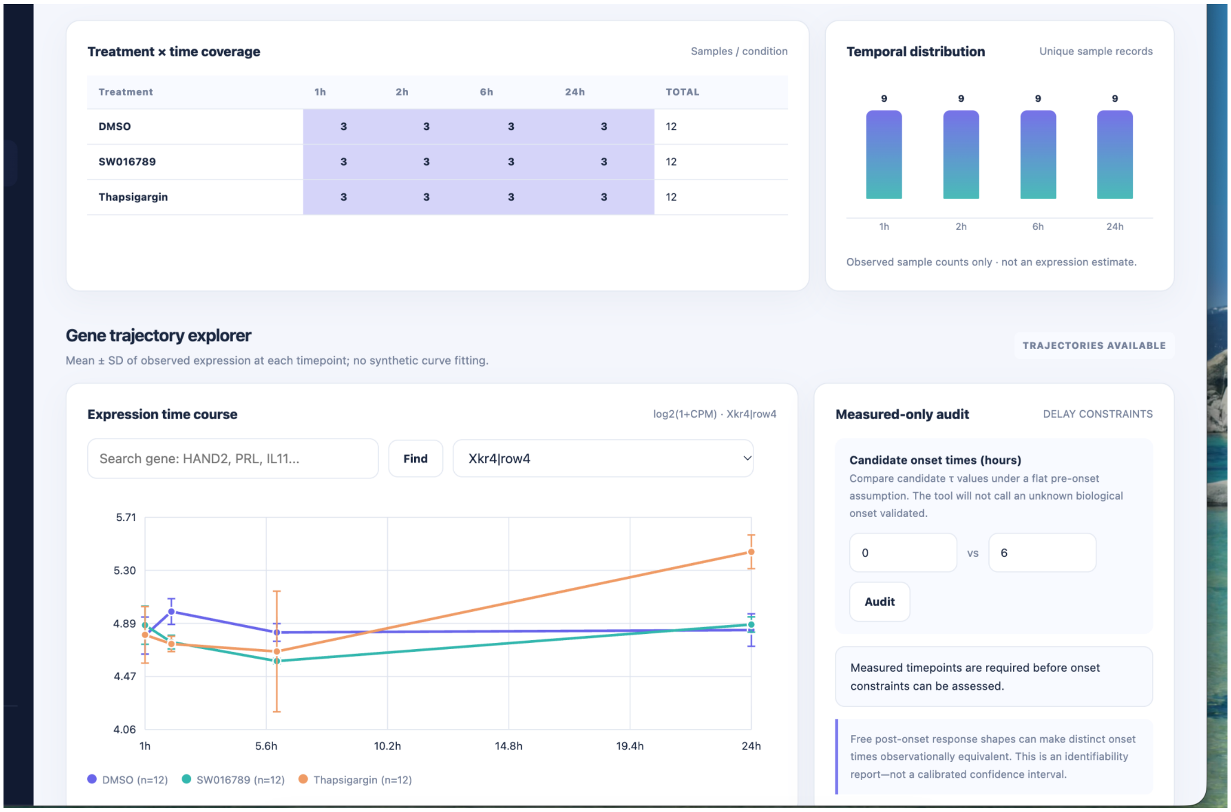


**Figure S1: Actual OnsetAudit screenshot, not an illustrative mock-up.** The locally running app displays imported GSE194200 treatment-by-hour coverage and measured RNA-seq trajectories for a mouse MIN6 cell-line experiment. This is unrelated to PGR knockdown in human endometrium and not a measurement of molecular-onset time.

OnsetAudit is a local Python application with an HTML/JavaScript user interface. It validates a time-course sample sheet and gene-expression matrix, presents observed trajectories, runs an experiment-grouped elapsed-exposure-time classification demonstration and exports a research report. The original GSE194200 demonstration uses two interventions (SW016789 and Thapsigargin) with DMSO vehicle controls. Among 24 active samples, 22 were both accepted and correctly assigned to one of the recorded elapsed-time categories, and among 12 distinct vehicle controls, 10 were rejected. These are within-study experimental-group holdouts, *not* independent cross-study validation, and no biological start time was independently labeled.

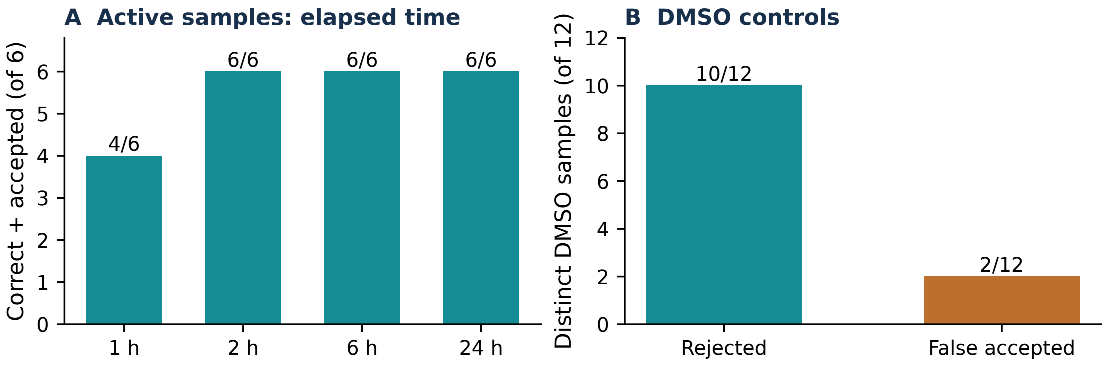


**Figure S2: Historical OnsetAudit elapsed-time demonstration.** A frozen GSE194200 report summarizes correctly timed active samples and rejected or incorrectly accepted DMSO controls. These numbers assess a technical workflow, not the human endometrial timing hypotheses in the main paper.

#### S3. Exact mathematical counterexamples and model boundary

For $f(t) = e^{kt}$, $f(t-\tau) = e^{-k\tau} f(t)$. For $f(t) = \alpha t + \beta$, $f(t-\tau) = f(t) - \alpha\tau$. For one fixed shift $\tau_0$, if $q$ is $\tau_0$-periodic, $f(t) = e^{kt} q(t)$ obeys $f(t - \tau_0) = e^{-k\tau_0} f(t)$. These are exact on the entire domain where the functions are specified, and not biological claims. For constant profiles, a shift is also not recoverable. Additional unknown onset phase can alias an additive shift by construction. Restricting amplitude to be positive changes Eq. (3) of the main text when the best signed affine coefficient is negative. The historical eight-gene V103 study investigated nonnegative amplitudes and modelled time-shift geometry separately, and should not be pooled with the all-feature analysis reported below.

The actual GSE246591 evaluation imposes $f(t<0) = f(0)$, unlike the globally defined exponential witness. These modeling choices are not interchangeable. The measured 2-hour value is absent from the GSE246591 data; generating it by interpolation would not add observational information. A future experiment needs an actual sample at the proposed time to improve identifiability.

##### S3.1. Feature eligibility and identifiable combinations

For a fixed candidate shift on a common finite time grid, suppose $\mathbf{f}_\tau = c_\tau \mathbf{f} + d_\tau \mathbf{1}$. Because adding a constant does not change the maximum-minus-minimum range and multiplying a vector scales the range by its absolute value, $R_\tau = |c_\tau| R_0$ exactly. Under the manuscript's two dynamic-range gates $R_0 \geq r$ and $R_\tau \geq r$, such an aliased feature is retained if and only if $R_0 \geq r$ and $|c_\tau| \geq r/R_0$. For a globally exponential function with growth constant $k$, this is $R_\tau = e^{-k\tau} R_0$. If $k > 0$, greater positive shifts can lead to exclusion; if $k < 0$ the range increases; an affine reference has $R_\tau = R_0$. This identity does *not* automatically hold under the empirical pre-zero clamping boundary, $f(\max(t-\tau, 0))$, even for an exponential. It concerns this particular model-dependent range screen, not the validity of all conventional independent filtering [1].

More generally, the least-squares geometric residual can never exceed the constant-only fit residual. By the range–variance inequality, $\delta_{\mathrm{RMS}} \leq \mathrm{range}(\mathbf{f}_\tau)/2$. For $R_0 > 0$, this gives $\delta_{\mathrm{RMS}}/R_0 \leq R_\tau/(2R_0)$. This is a deterministic upper bound, not an estimate of the probability of an onset error. It explains why candidates excluded for small $R_\tau$ merit separate reporting rather than silent deletion; it does not guarantee that any particular excluded feature passes the manuscript's 0.05 near-zero cutoff.

For the unrestricted exponential $y(t) = b + a\exp[k(t-\tau)]$, the identifiable combination is $A = a\exp(-k\tau)$. With nonzero $A$, nonzero $k$ and three equally spaced distinct observation times, the finite difference ratio is $e^{kh}$, giving the finite-difference rate identity stated in the main paper. If $A = 0$ or $k = 0$, this ratio is undefined; noise or weak curvature can make rate identification unstable. The shift derivative is confounded with the amplitude derivative, whereas the rate derivative need not be; this is a standard example of identifiable parameter combinations [2].

##### S3.2. Retrospective shift-only exclusion audit

The independent script `scripts/p7_selection_identifiability_audit.py` uses the already archived, per-feature $\delta$ results without refitting or changing the primary analysis. For each culture and candidate shift it reports (i) the number passing the original $R_0 \geq 0.5$ gate, (ii) additional exclusions from the modeled $R_\tau \geq 0.5$ gate, (iii) the near-zero residual fraction in both retained and excluded sets, and (iv) a fixed-feature intersection across all four shifts. The selection-audit script is supplied, but its numerical output is not included until it has been executed against the archived per-feature CSV and the source hashes verified. No selection-effect direction is claimed here.

#### S4. Full measured-source GSE246591 results

**Table S1:** All 16 author-run summaries of model-conditional projection geometry for the original count matrix (60,676 source features). Medians and quantiles are over eligible feature rows separately for each culture and shift. The last column is the descriptive fraction below 0.05, *not* a posterior probability or statistical error rate.

| Culture | $\tau$ h | Features | Median $\delta_{\mathrm{RMS}}$ | Median normalized | q10–q90 normalized | $< 0.05$ |
|---|---|---|---|---|---|---|
| S169 | 3 | 14019 | .2023 | .2203 | .1164–.2840 | .0058 |
| S169 | 6 | 11746 | .1988 | .2057 | .1182–.2922 | .0020 |
| S169 | 12 | 10931 | .2239 | .2259 | .1305–.3381 | .0002 |
| S169 | 24 | 9116 | .2361 | .2186 | .1233–.3540 | .0005 |
| S170 | 3 | 13694 | .1477 | .1534 | .0833–.2433 | .0205 |
| S170 | 6 | 13584 | .1965 | .2143 | .1218–.2950 | .0037 |
| S170 | 12 | 12173 | .2185 | .2261 | .1319–.3298 | .0018 |
| S170 | 24 | 10281 | .2218 | .2088 | .1214–.3310 | .0032 |
| S506 | 3 | 11422 | .1380 | .1589 | .0770–.2494 | .0320 |
| S506 | 6 | 11254 | .1778 | .2099 | .1062–.3019 | .0112 |
| S506 | 12 | 10300 | .2025 | .2276 | .1237–.3438 | .0044 |
| S506 | 24 | 8225 | .2089 | .2130 | .1169–.3380 | .0072 |
| S508 | 3 | 13114 | .1510 | .1649 | .0858–.2528 | .0188 |
| S508 | 6 | 12775 | .1856 | .2073 | .1119–.3007 | .0064 |
| S508 | 12 | 11629 | .2117 | .2307 | .1285–.3409 | .0023 |
| S508 | 24 | 9690 | .2150 | .2131 | .1240–.3310 | .0026 |

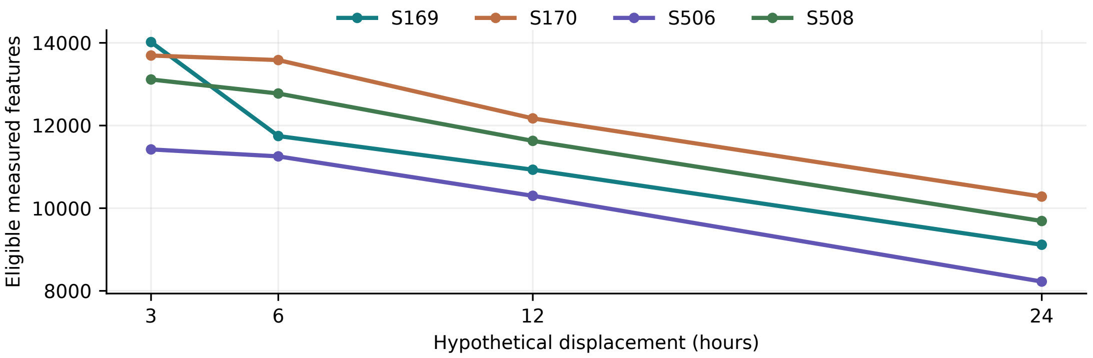


**Figure S3: Shift-dependent number of eligible features.** Larger shifts often cause more feature exclusions under the stated dynamic-range gates. Hence differences in the main-text median curves do not compare an invariant gene set.

#### S4.1. Separating retained signal and affine shape separation

For a measured baseline curve $\mathbf{f}$ and its model-implied displacement $\mathbf{f}_\tau$, the final descriptive audit computes $R = \text{range}(\mathbf{f}_\tau)/\,\text{range}(\mathbf{f})$ and $S = \|(I - P_X)\mathbf{f}_\tau\|_2/\|\mathbf{f}_\tau - \overline{f_\tau}\mathbf{1}\|_2$. The shifted profile uses the same ten recorded times, piecewise-linear interpolation, and constant pre-zero extension as the main analysis. When the shifted profile is constant, $S$ is *undefined*, not zero. For both nonconstant profiles, $S = \sqrt{1 - r^2}$ with $r$ the Pearson correlation under an unconstrained *signed* fitted amplitude. The joint summary is not a noise-calibrated delay classifier.

**Table S2:** Two distinct sources of small projection residuals ($D < 0.05$) using only user-reported aggregated results. Columns $R$ and $S$ are *medians among near-zero* feature records, not medians over the entire stratum. At 3 h the stratum includes initially dynamic features; at 24 h it is split by candidate-shift eligibility. The 0.05 cutoff is descriptive. The corresponding feature-level source was analyzed on the author's Mac.

| Culture | Shift | Stratum | Near-zero / total | Median $R$ | Median $S$ |
|---|---|---|---:|---:|---:|
| S169 | 3h | Initial | 81/14,766 | .975 | .138 |
| S170 | 3h | Initial | 284/14,871 | .971 | .139 |
| S506 | 3h | Initial | 365/12,745 | .968 | .131 |
| S508 | 3h | Initial | 252/14,606 | .968 | .141 |
| S169 | 24h | Excluded | 322/5,650 | .136 | .989 |
| S170 | 24h | Excluded | 108/4,590 | .153 | .969 |
| S506 | 24h | Excluded | 114/4,520 | .186 | .884 |
| S508 | 24h | Excluded | 186/4,916 | .173 | .865 |
| S169 | 24h | Retained | 5/9,116 | .578 | .228 |
| S170 | 24h | Retained | 33/10,281 | .616 | .236 |
| S506 | 24h | Retained | 59/8,225 | .582 | .234 |
| S508 | 24h | Retained | 25/9,690 | .600 | .225 |

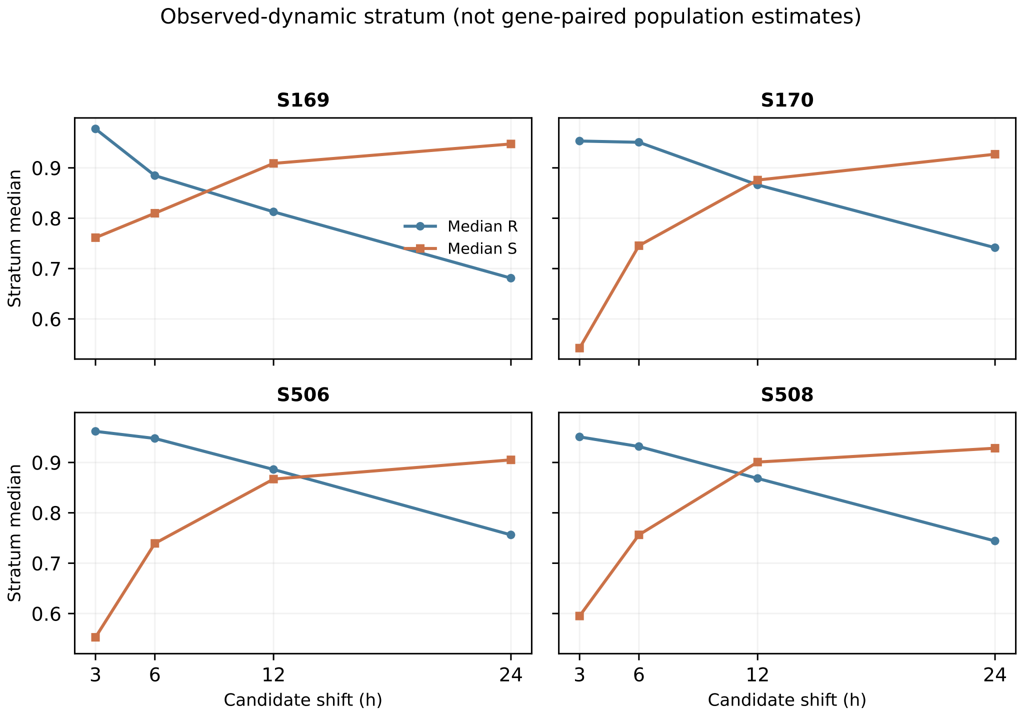


**Figure S4: Median two-component diagnostics over the initial observed-dynamic stratum.** These descriptive culture medians are computed separately for each candidate shift. Feature identities recur across candidate shifts, and the modeled shifted values are not independent observations. This figure does not show joint feature-level $(R, S)$ scatter.

Among initially dynamic culture–feature records (56,988 across the four cultures), the additional shifted-range gate at 24h excluded 19,676 (34.5%). Within the excluded, 730 were near-zero by $D < 0.05$; 557/730 (76.3%) had $R < 0.2$, 20/730 (2.7%) had $S < 0.2$, and one exactly flat shifted profile had undefined $S$. At 3h, among 982 near-zero initially dynamic records, 977 (99.5%) had $S < 0.2$ while none had $R < 0.2$. These two exploratory cutoffs were chosen for display and are *not* statistically calibrated. Ratios and percentages pool dependent feature records; no confidence intervals, hypothesis tests, independent donor replication or biological-onset inference are claimed.

The retrospective comparison is reproduced by `scripts/p7_dual_component_audit.py` using the same deposited counts checksum as the primary analysis. Because the author-run per-feature audit ZIP was not transferred into this Overleaf archive, the present figure is drawn *only from reported aggregate summaries*, not a replacement for a feature-level replication or a fabricated point cloud. To reproduce all individual points, rerun the bundled script with the full GSE246591 counts and prior per-feature projection export.

### S5. Exploratory reference-shape sensitivity

A retrospective panel of 48 features was selected for substantial measured range in every culture, using all ten time points, so the feature selection was *not nested inside validation.* Three approximation families were compared: linear interpolation, a Gaussian process with fixed 12-hour radial-basis length scale and regularization 0.08, and a grid search of two-sigmoid "impulse-like" shapes. These are neither the official implementations nor equivalent targets of DEtime [3] and ImpulseDE2 [4]. The leave-one-time-point-out (LOO) measure omits each measured time in turn and computes median per-feature root-mean-square reconstruction error (RMSE) across all ten held-out times. Endpoints require extrapolation. The shape diagnostic then recalculates the three-hour signed-amplitude projection on each fitted curve.

**Table S3:** Retrospective 48-feature model-shape pilot. First three numeric columns are median LOO RMSE in $\log_2(1+\text{CPM})$. Last three are the three-hour median normalized projection residuals; this is an *exploratory numerical sensitivity table only.*

| | LOO reconstruction error | | | 3h normalized residual | | |
|---|---|---|---|---|---|---|
| Culture | Linear | Fixed GP | Impulse-like | Linear | Fixed GP | Impulse-like |
| S169 | .6583 | 1.1395 | .8695 | .0945 | .0426 | .0345 |
| S170 | .6239 | 1.1573 | .8530 | .0802 | .0444 | .0350 |
| S506 | .7366 | 1.2696 | .9399 | .0569 | .0299 | .0271 |
| S508 | .5946 | 1.1762 | .7151 | .0677 | .0372 | .0333 |

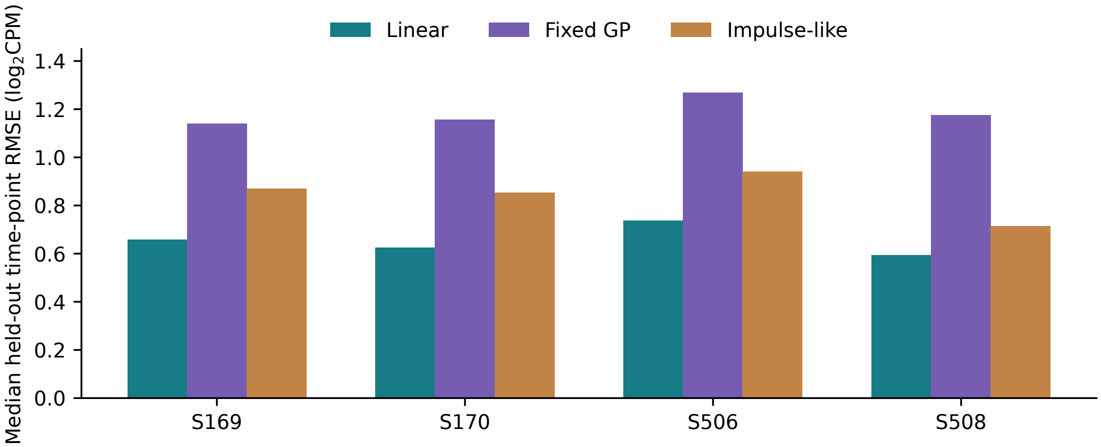


**Figure S5: Retrospective reference-shape reconstruction.** Retrospective reference-shape LOO reconstruction. Smaller values indicate better reconstruction *for this fixed pilot setup*, not algorithmic superiority or performance on held-out genes.

**Numerical warning.** The original runtime printed overflow, division-by-zero and invalid-operation warnings during matrix calculations for the GP/impulse approximations. A subsequent export check found 576/576 finite LOO rows, 48/48 finite shape-diagnostic rows and 12/12 finite summary rows; that confirms exported completeness only and *does not* prove that non-finite intermediate calculations had no impact. Accordingly, the above results are retained for transparent sensitivity reporting and are *not* used to claim validated model ranking, optimized hyperparameters, calibrated uncertainty or superiority over the published DEtime and ImpulseDE2 implementations.

### S6. Biological context: keep estimands separate

**Table S4:** Distinct inferential analyses of the GSE51981 frozen PGR-dependency score. The MSE comparison includes 28 endometriosis and eight strict-control samples. The phase model was fitted on the larger phase-stratified analysis set ($n = 109$). CI means 95% confidence interval; HC3 denotes the heteroskedasticity-robust covariance specification.

| Model / estimand | Difference | CI lower | CI upper | $p$ |
|---|---|---|---|---|
| Phase-interaction HC3, MSE effect | −.154922 | −.222227 | −.087617 | $1.3860 \times 10^{-5}$ |
| MSE-only HC3, unadjusted | −.154922 | −.223889 | −.085955 | $6.2474 \times 10^{-5}$ |
| MSE-only HC3, three composition proxies | −.076561 | −.133443 | −.019678 | .009975 |

An earlier MSE-only Welch test yielded $p \simeq 4.51 \times 10^{-5}$; its p-value cannot be paired with the phase-model HC3 confidence interval. The registered GSE51981 abstract interval comes from the *phase-model* HC3 row rounded to three decimals. For GSE6364, the frozen 25 October v0.24 primary experiment used 17 MSE biopsies (nine endometriosis, eight controls), the original $\log_2$(GEO-supplied GCOS signal) representation, and source weights carried over without new target-gene selection. The Welch comparison was $-0.032765$ (95% CI $[-0.067656, +0.002126]$, $p = 0.0637523$). Its quantile-normalization sensitivity yielded $-0.027399$ (95% CI $[-0.060041, +0.005243]$, $p = 0.0934664$). These are score differences on a separate clinical cohort, not observations of delay. Both original human datasets use array measurements, but their source transformations and target populations must not be pooled.

### S7. Reproducibility and scope of claims

The main paper is self-contained. This Overleaf archive includes editable LaTeX, the BibTeX bibliography, publication figures, the measured GSE246591 summary transcribed from the author's 40-sample audited run, input-file SHA-256, and the real OnsetAudit screenshot. For a full independent rerun, download the original GEO GSE246591 source and execute `scripts/compute_gse246591_delta.py` on a machine with Python, NumPy and pandas. Inspect its SHA-256 and the complete per-feature output; the 16-row summary in this archive is not a replacement for the source matrix. The visible source repo is `https://github.com/Helen2813/paper7`; README and exact commit must be checked before submission. The original OnsetAudit archived report is included in `evidence/`. Model limits are: no verified true lag, no variance-calibrated delay CI, no matched official DEtime/ImpulseDE2 benchmark, no independently validated drug-identity model, and no evidence of causal PGR timing change.
